# The cellular architecture of wheat leaves supports a conservation of spatial patterning of the mesophyll in grasses

**DOI:** 10.1101/2025.03.16.643523

**Authors:** Emma White, Matthew J. Wilson, Richard Summers, Cristobal Uauy, Andrew Fleming

**Affiliations:** School of Biosciences, University of Sheffield, Western Bank, Sheffield S10 2TN, United Kingdom; RAGT Seeds Limited Grange Road, Ickleton, Essex CB10 1TA, United Kingdom; John Innes Centre, Norwich Research Park, Norwich NR4 7UH, United Kingdom

**Keywords:** Cell Shape, Cell Size, Mesophyll, Wheat, Triticum, Leaf Development

## Abstract

Classical botany identifies a clear adaxial/abaxial (palisade/spongy) patterning of the mesophyll in leaves of eudicots but suggest that there is no comparable overt pattern within the mesophyll of most grasses. In this paper, we report on an analysis of leaves from across ploidy levels of Triticum species which reveals a patterning of mesophyll cellular architecture that has been maintained during the evolution/selection of wheat. Taking a quantitative approach to the analysis of 2D and 3D images, we show that upper and lower mesophyll cells are larger than the middle mesophyll cells, which are themselves more lobed than the mesophyll cells adjoining the upper and lower epidermis. This mesophyll patterning is superimposed on a gradual increase of cell size with ploidy, suggesting that it is evolutionarily ancient. Taken in the context of data revealing patterning of the rice mesophyll, our data support the hypothesis that grass leaves do display distinct cellular patterning across the adaxial/abaxial axis, the functional significance of which awaits full elucidation.

## INTRODUCTION

Mesophyll cells constitute the vast bulk of tissue in most leaves. They are the primary site of photosynthesis and, at an organ level, display numerous adaptations to optimise the capture of light and CO_2_ required for carbon fixation/assimilation, while at the same time limiting the loss of water. These adaptations, which can be physiological or structural in nature, can be highly species specific and, moreover, change in response to environmental conditions. Nevertheless, there are general patterns of mesophyll structure which reflect major sub-divisions in plant evolution. For example, eudicot leaves are traditionally viewed as being composed of two distinct layers (palisade and spongy), with the constituent mesophyll cells displaying distinct shapes and patterns of cell separation (Esau, 1965; Pyke, 2012). This distinct cellular architecture is thought to reflect primary functions of the layers in, e.g., light absorption and gas exchange (Flexas *et al.,* 2012; Evans *et al.,* 2009; Vogelmann & Martin, 1993), although, in reality, there is a gradient of function across the leaf adaxial/abaxial gradient which is rarely absolutely tightly limited to such spatial boundaries.

This classical definition of palisade/spongy mesophyll across the eudicot leaf adaxial/abaxial contrasts with most grass leaves where mesophyll architecture appears much more uniform (Chonan, 1978). Our recent work indicated that rice leaves do not have a uniform mesophyll architecture and that cells located towards the adaxial or abaxial sides of the leaf can be distinguished via quantitative analysis of cell size and shape (Sloan *et al.,* 2023). We speculated that this distribution of mesophyll cell size and shape might influence physiological and/or mechanical parameters of leaf function. Since our analysis was restricted to rice, our previous investigation led to the question of whether our observations were restricted to this species or reflected a more general situation in grass leaves. We therefore turned our attention to wheat, another grass species of significant agronomic interest and which also presents a well characterised array of species/cultivars across evolutionary space for analysis.

Modern bread wheat (*Triticum aestivum*) is a hexaploid derived from tetraploid and diploid progenitors. Around 400,000 years ago, wild diploid *T. uratu* (2n = 2x = 14, genome A^u^A^u^) hybridised with the B genome ancestor of modern wheat (*Aegilops speltoides,* 2n = 2x = 14, genome BB) to produce wild tetraploid emmer wheat (*T. turgidum ssp. dicoccoides*, 2n = 4x = 28, genome A^u^A^u^BB) (Dvorak *et al.,* 2012). Wild emmer began to be cultivated around 10,000 years ago to create a cultivated emmer (*T. turgidum ssp. dicoccum,* 2n = 4x = 28, genome A^u^A^u^BB) (Feldman & Kislev, 2007; Kislev *et al.,* 1992). After another 1000 years, the cultivated emmer spontaneously hybridized with another goat grass, similar to the B genome progenitor (*A. tauschii,* 2n = 2x = 14, genome DD) to produce an early spelt variety (*T. spelta*, 2n = 6x = 42, genome A^u^A^u^BBDD). However, the idea that *T. spelta* is the ancestor of modern bread wheat has been disputed (Dvorak *et al.,* 2006), and instead the idea presented that cultivated wheat ancestry is more complicated, involving factors such as gene flow from wild cereals (Dvorak *et al.,* 2011). Irrespective of the continuing debate on the precise character of wheat evolution, the extensive research in this area means that we have access to a range of related Triticum species of varying ploidy and cultivation status which can be used to address questions related to leaf structure/function over evolutionary time.

Classical investigations of wheat leaf structure indicated an overall structure comparable to other grasses in that there was no overt separation of mesophyll into adaxial or abaxial layers, and instead, similar to other monocots, leaf cells have distinctly lobed shapes and are patterned across the proximal-distal axis, interspaced by major and minor veins (Jellings & Leech, 1984; Parker & Ford, 1982). These studies utilised traditional microscopy approaches looking at embedded sections of tissue in 2D. In the intervening time, elegant and powerful confocal microscopy-based methods have been developed which, combined with advanced image processing approaches, allow quantification of cell size, shape and arrangement in 3D. These approaches have been applied to the wheat leaf (Wilson *et al.,* 2021) but have not been used to investigate the potential for mesophyll patterns across the adaxial/abaxial axis.

In this paper, we report on an investigation of mesophyll size and shape distribution in a range of extant Triticum species encompassing the spectrum of wheat evolution and cultivation. Using a combination of quantitative approaches, we show that wheat leaves have a structured mesophyll, supporting the hypothesis that grass leaves have a cellular architecture that is spatially more complex than previously thought.

## METHODS

### Plant Material and Growth

The plant materials used in this study can be found in **Table 1**.

**Table 1.** Triticum lines used in this study. Lines are referred to by the simplified ‘Name’ in the text, and colour coded by ploidy and cultivation status. The lines used for confocal imaging were a subset of the total panel: 2n – TRI6735; 4nW – TRI18505; Emmer – TRI16877; Durum – Voilur; 6n – Fielder.

| Ploidy | Species | Line Code | Status | Name |
| --- | --- | --- | --- | --- |
| Diploid | *Triticum baeoticum* | TRI 18344 | Wild | *T. baeoticum* |
| Diploid | *Triticum uratu* | TRI 6735 | Wild | *T. uratu* |
| Tetraploid | *Triticum turgidum* (subsp. *dicoccoides*) | TRI18505 | Wild | *T. dicoccoides* |
| Tetraploid | *Triticum turgidum* (subsp. *dicoccoides*) | TRI18530 | Wild | *T. dicoccoides* |
| Tetraploid | *Triticum araraticum* | TRI16599 | Wild | *T. araraticum* |
| Tetraploid | *Triticum araraticum* | TRI18513 | Wild | *T. araraticum* |
| Tetraploid | *Triticum turgidum* (subsp. *dicoccon*) | TRI16877 | Domesticated (Landrace) | Emmer |
| Tetraploid | *Triticum turgidum* (subsp. *dicoccon*) | TRI28049 | Domesticated (Landrace) | Emmer |
| Tetraploid | *Triticum turgidum* (subsp. *dicoccon*) | TRI14734 | Domesticated (Landrace) | Emmer |
| Tetraploid | *Triticum durum* (subsp. *durum*) | VOILUR | Domesticated (Modern) | Durum |
| Tetraploid | *Triticum durum* (subsp. *durum*) | AVENTADUR | Domesticated (Modern) | Durum |
| Tetraploid | *Triticum durum* (subsp. *durum*) | ANVERGUR | Domesticated (Modern) | Durum |
| Hexaploid | *Triticum aestivum* | COUGAR | Domesticated (Modern) | *T. aestivum* |
| Hexaploid | *Triticum aestivum* | FIELDER | Domesticated (Modern) | *T. aestivum* |

Plants were sown in F2+S compost (ICL, Suffolk, UK) and placed into a controlled environment growth chamber (Conviron PGR15; Conviron, Winnipeg, MB, Canada) (air temperature = 21°C:16°C day:night, 16h photoperiod = 400 μmol m^-2^ s^-1^, 8h dark, 60% relative humidity, ambient CO_2_). After germination and 1 week of growth, the seedlings were transplanted into larger pots containing 6:1 mix of F2+S compost and Perlite (Sinclair Pro, Cheshire, UK), and five grams of Osmocote Exact 5-6 slow-release fertiliser (ICL, Suffolk, UK) was added to each pot and mixed into the top layer of compost. Plants were watered three times per week throughout the 28-day growth period. Leaf width was measured at the broadest part of the 5^th^ leaf from the main tiller at 28 days after sowing.

### 2D Mesophyll Imaging

After 28 days of plant growth from sowing, three sections approximately 1 cm in length were taken from approximately one third of the way down from the tip of the fifth leaf on the main tiller of each plant and immediately placed into a fixative comprising of 3:1 ethanol:acetic anhydride (v/v) in glass vials. The samples were placed into a vacuum pump for one hour, before being stored at room temperature for at least 48 hours. Samples were then washed with a 50% ethanol solution for 15 minutes, before being stored in 70% ethanol at 4°C.

Samples for Technovit® (TAAB Laboratories Equipment Ltd, UK) sectioning were transferred to 1:1 EtOH:Technovit® 7100 base liquid and vacuum infiltrated for 1 hour before being left for 24 hours at room temperature. The samples were then moved to 100% Technovit® base liquid and stored at 4°C for 48 hours, before being transferred to Technovit® 1 (1 g Hardener 1 per 100 ml Technovit® base liquid), and stored at 4°C for a further 24 hours.

Samples were then stained for 5 minutes using neutral red dye and embedded in Technovit® 3040 resin and were sectioned at 8 μm using a Leica Microtome. Six transverse and six longitudinal sections were taken from each line and then stained with toluidine blue + 0.05% Borax before being mounted onto microscope slides using Eukitt® Quick-hardening mounting medium (Sigma-Aldrich, UK).

Images were taken on an Olympus BX51 microscope with an Olympus DP71 camera and Cell B imaging software using the 20x objective. Two images from each section were taken where possible, resulting in a maximum of 12 images per plant for 4-6 plants per line. Transverse sections were taken between the 2nd and 3rd veins out from the midvein.

### 2D Mesophyll Cell Measurements

Each mesophyll cell was allocated a ‘layer’ identity depending on its location in the cell (See **Figure 1**). ‘Upper’ layer mesophyll cells were identified as being directly adjacent to the adaxial epidermis, ‘Lower’ mesophyll cells being directly adjacent to the abaxial epidermis, ‘Radial’ mesophyll cells being directly adjacent to bundle sheath cells surrounding the vasculature and ‘Middle’ mesophyll cells being only directly adjacent to other mesophyll cells. Mesophyll cell area and shape was measured by digitally drawing around each cell using images of both transverse and longitudinal sections in FIJI (ImageJ 5.3g; Schindelin *et al*., 2012) using an in-house macro (.ijm). Every cell within was outlined by hand, and area (μm^2^), perimeter (μm), circularity, cell length (Feret), cell width (MinFeret), and convex hull perimeter (μm) measurements were taken. Mesophyll cell lobing area was calculated as convex hull area minus cell area. For transverse sections, circularity was used to quantify shape and lobing area for longitudinal sections.

**Figure 1.**
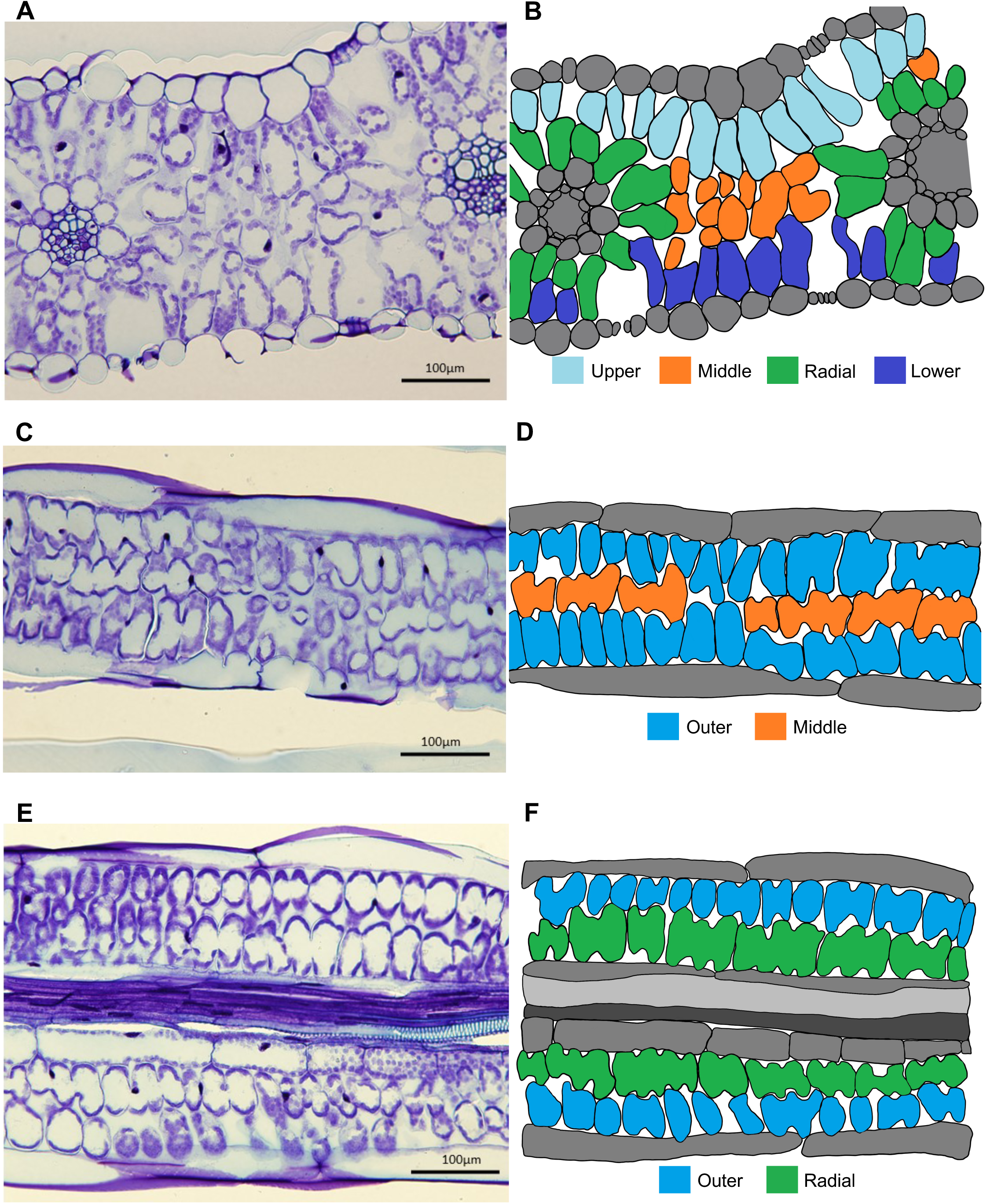
The assignation of mesophyll cells to layers in a typical wheat leaf. **A)** Transverse section of a durum leaf (5^th^ leaf at 28 days after sowing); **B)** Diagram of A in which cells have been assigned to spatial location, as indicated by the colour code (Upper = upper (adaxial) mesophyll; Middle = middle mesophyll; Radial = arranged radially adjacent to a vascular bundle; Lower = lower (abaxial) mesophyll); **C)** Longitudinal section of a durum leaf outside a plane containing no radial cells; **D)** Diagram of C, colour coded as in B; **E)** Longitudinal section of a durum leaf showing radial cells; **F)** Diagram of E, colour coded as in B.

The data from all ImageJ macros used in this project were exported as an annotated image, a zipped folder containing ROI data and CSV files. The numerical data was then organised and analysed using RStudio (ver. 4.1.0). The script outputs data for variables such as cell area and circularity/lobing for each leaf.

### 3D Imaging

Leaf sections were treated with chloroform for 10 minutes to remove leaf cuticular waxes (Wuyts *et al.,* 2010), then transferred to 70% ethanol for 15 minutes before rinsing in 50% ethanol and finally in dH_2_O. The samples were then suspended in 0.2 M NaOH solution containing 1% Sodium dodecyl sulphate (w/v) (SDS) for 15 minutes before rinsing three times with distilled water. The leaf tissue was then treated with a starch digester solution consisting of phosphate buffered saline (PBS), 0.1% Tween-20 and 0.01% (w/v) alpha-amylase (Sigma-Aldrich, UK) and incubated at 37°C overnight.

After incubation, samples were treated with a freshly prepared solution of 1% (v/v) periodic acid (Sigma-Aldrich, UK) for 40 minutes, before rinsing in dH_2_O. Samples were then stained using propidium iodide (PI, Sigma-Aldrich, UK). Leaf sections were treated in a Pseudo-Schiff PI solution consisting of 0.1 M Na_2_S_2_O_5_, 0.15 N HCl, 0.01% PI for 4 hours in dark conditions. After the staining period, samples were washed three times in water and left overnight in the third rinse at 4°C.

The samples were then cleared in a solution consisting of 200 g chloral hydrate (Sigma-Aldrich, UK), 20 ml glycerol and 30 ml dH_2_O for 6 hours. Once cleared, the leaf sections were blotted using a tissue to remove excess liquid. Each section was then dissected along the midrib to allow for the preparation of flatter samples. One piece of the dissected sample was flipped so that the adaxial side was facing up and was placed directly onto coverslip using a few drops of mountant consisting of: 3 g 20% (v/v) Arabic gum, 10 g chloral hydrate and 1 g glycerol and mounted onto microscope slides.

The leaf sections were imaged using a ZEISS Airyscan confocal microscope. The PI stain was excited using the 561 nm DPSS Diode laser. Scans were performed at a resolution of 1024 x 1024 pixels, with a pixel dwell of 1.06 μs/pixel and with a z-step interval of 0.3 μm per slice. The 20x dry objective was used. Once an appropriate area was chosen based on sufficient absence of noise, vasculature and significant substomatal cavities, the capture area was zoomed to 2x before beginning the experiment. Correction was used to increase the laser power to manually inputted values at regular intervals throughout the z-stack, between a laser power of roughly 3-18%.

### 3D Image Processing and Analysis

3D reconstruction and segmentation of the leaf mesophyll cells was carried out using MorphographX software (Barbier de Reuille *et al.,* 2015/Strauss *et al.,* 2022). Stacks were initially converted into TIFF format using FIJI/ImageJ. The TIFF image files were loaded in MorphoGraphX and the built-in, pre-trained CNN for cell boundary prediction using the process “Stack/CNN/UNet3D” (Vijayan *et al.,* 2021) was used prior to segmentation. The correct voxel size was inputted, and gaussian blur was added to the stack at a value of between 0.3 and 0.7 in all directions, optimised for each individual sample.

Segmentation of the sample was the carried out using the ITK Threshold Autoseeded Watershed function, set to a threshold of 1000-1500, manually optimised for each sample. This value was altered appropriately depending on whether the stack was over-or under-segmented. The seeding was examined, and labels were fused and reseeded where necessary. Any under-segmented regions and intercellular airspaces were removed at this stage.

The mesh was generated using a 3D marching cubes algorithm, with a cube spacing of 1μm and three smooth passes. Once this mesh was created, final cleaning up of the sample was carried out so that all incomplete cells touching the edges of the z-stack were removed. Any remaining artefacts were also removed along with other any non-mesophyll cells remaining. Data relating to cell volume and surface area were exported as spreadsheets and heatmaps, and the number of ‘lobes’ per each cell was manually documented.

Prior to generation of heat-maps, the mesh was rescaled by 1.55x in the z-direction to compensate for any spherical aberration and axial distortion experienced during imaging due to differences in refractive index, as described by Diel *et al*. (2020).

### Data Analysis

Data recorded and outputted from ImageJ (2D) and MorphographX (3D) was statistically analysed and graphed using GraphPad Prism 9 (ver. 9.3.1; Graphpad Software Inc.). All error bars on graphs indicate one standard deviation (SD) from the mean.

## RESULTS

### Analysis of 2D mesophyll cell size and shape in wheat reveals patterning across the adaxial-abaxial leaf axis

Observation of both cross-sections (**Figure 1A**) and longitudinal sections of (**Figure 1C,E**) of wheat leaves suggested that mesophyll cells can be assigned to layers, as highlighted by the relevant diagrams (**Figure 1B,D,E**). Thus, in cross-sections cells can be assigned to “Upper”, “Middle”, “Radial” and “Lower” mesophyll (**Figure 1B**). In longitudinal sections outside of the vascular bundles the radial cells are not apparent (**Figure 1C**). Mesophyll cells in these sections can easily be assigned to “Outer” layers (sub-adjacent to epidermis) and “Middle” layer (**Figure 1D**), but due to sample movement during processing it is difficult to confidently assign the “Outer” layers to either “Upper” or “Lower” sides of the leaf. Similarly, in longitudinal sections along vascular bundles (**Figure 1E**), although “Radial” cells could be easily designated (**Figure 1F**), tracking which of the “Outer” layers was upper or lower mesophyll was challenging, leading us to assign these cells simply to the “Outer” category (**Figure 1F**). The images shown in **Figure 1** relate to durum wheat (a cultivated tetraploid line) but similar apparent patterns and, therefore, cell assignations could be made in diploid wheat, other tetraploid lines (both cultivated and non-cultivated), as well as modern hexaploid bread wheat. The lines used for analysis in this investigation, with an indication of ploidy level, cultivation status and accession ID, are listed in **Table 1**.

To explore the apparent patterning across the leaf adaxial-abaxial axis, we performed a quantitative analysis of cell size and shape of cells in the different layers for plants of different ploidy and cultivation status. This was carried out using analyses of both leaf sections (2D analysis) (**Figures 2-4**) and of confocal-imaged and segmented datasets (3D analysis) (**Figure 5**).

**Figure 2.**
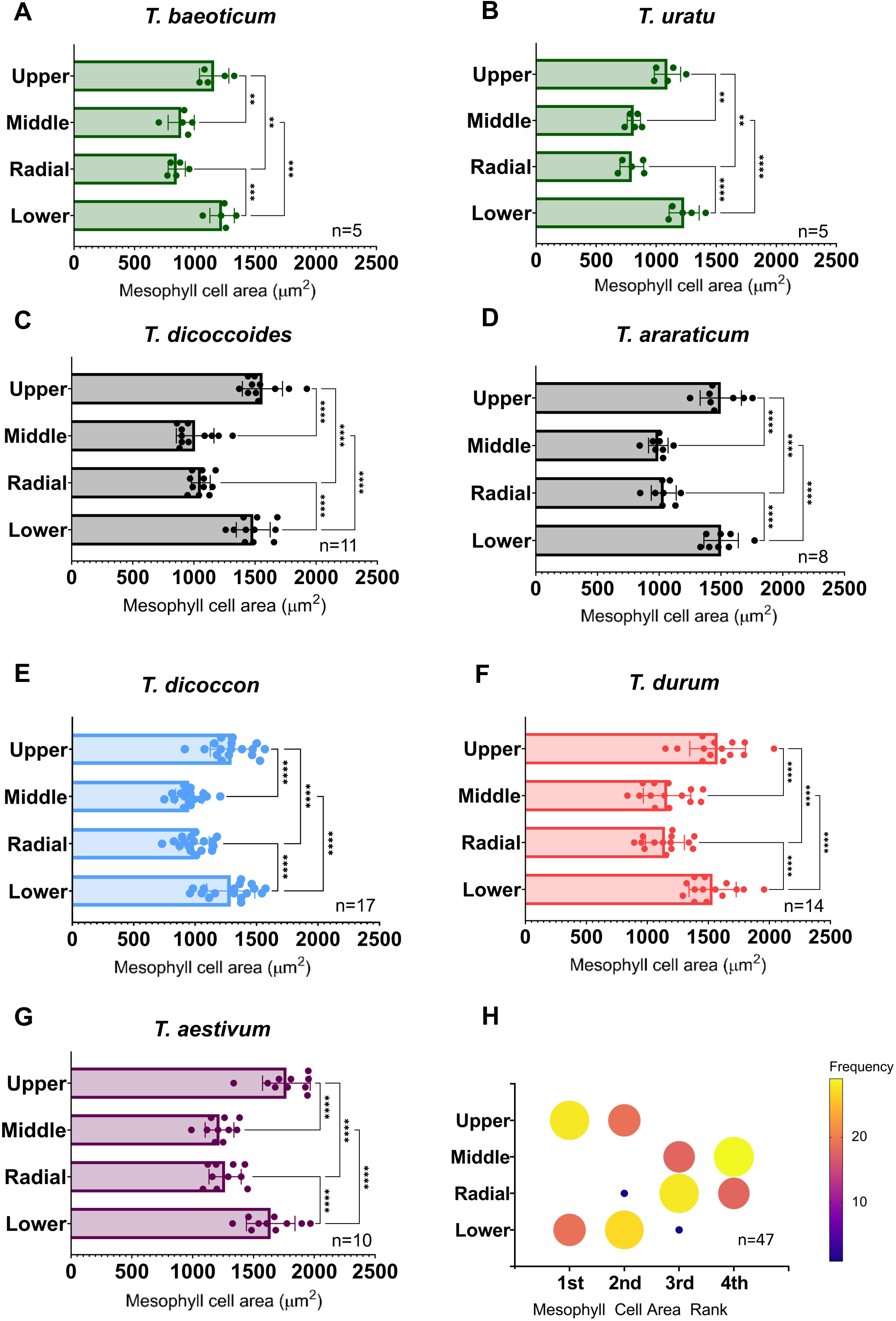
Analysis of transverse mesophyll cell area shows layers can be distinguished by size in diploid, tetraploid and hexaploid Triticum lines. **A-B**) Transverse area of cells in designated mesophyll layers of diploid species, *T.baeoticum* and *T.uratu* (as indicated); **C-D**) tetraploid species, *T. dicoccoides* and *T. araraticum*, as indicated; **E-F**) cultivated tetraploid species, *T. dicoccon* and *T. durum*, as indicated; **G**) hexaploid species, *T. aestivum*; **H**) Ranked data of transverse mesophyll cell area from all lines, separated by layer. Frequency of rank is indicated by colour and size of circle. Ranking is determined based on the frequency that cells in a particular layer rank 1st (largest) –4th (smallest) in terms of cell area. In A-G, error bars represent standard deviation from the mean. ANOVA and post-hoc Tukey tests were carried out to test for significant interactions between cell layers within each species/subspecies: p= <0.005, p= <0.0005, p= <0.0001. Statistical significance for ranked data in (H) was assessed using a Friedman test.

**Figure 3.**
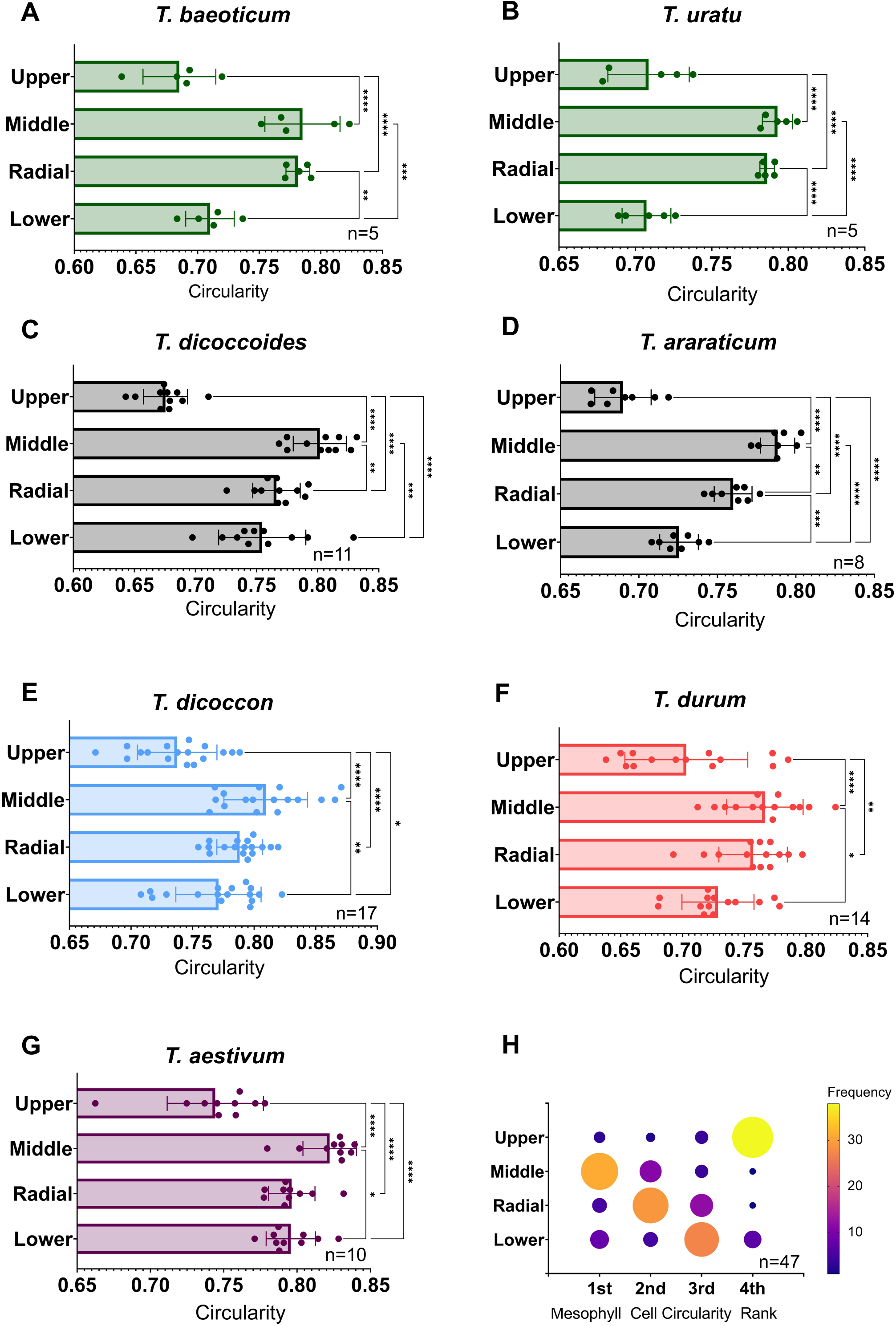
Analysis of transverse mesophyll cell circularity shows layers can be distinguished by shape in diploid, tetraploid and hexaploid Triticum lines. **A-B**) Transverse circularity of cells in designated mesophyll layers in diploid species, *T.baeoticum* and *T.uratu* (as indicated);. **C,D**) wild tetraploid *species T. dicoccoides* and *T. araraticum*, as indicated. **E-F**) cultivated tetraploid species, *T. dicoccon* and *T. durum*, as indicated **G**) hexaploid species, *T. aestivum*. **H)** Ranked data analysis of mesophyll cell circularity from all lines, separated by layer. Frequency of rank is indicated by colour and size of circle. Ranking is determined based on the frequency that cells in a particular layer rank 1st (most circular) –4th (least circular) in terms of cell circularity. Error bars represent standard deviation from the mean. ANOVA and post-hoc Tukey tests were carried out to test for significant interactions between cell layers within each species/subspecies: p= <0.05, p= <0.005, p= <0.0005, p= <0.0001. Statistical significance for ranked data in (H) was assessed using a Friedman test. Circularity represents a shape’s deviation from a perfect circle which has a value of 1.

**Figure 4.**
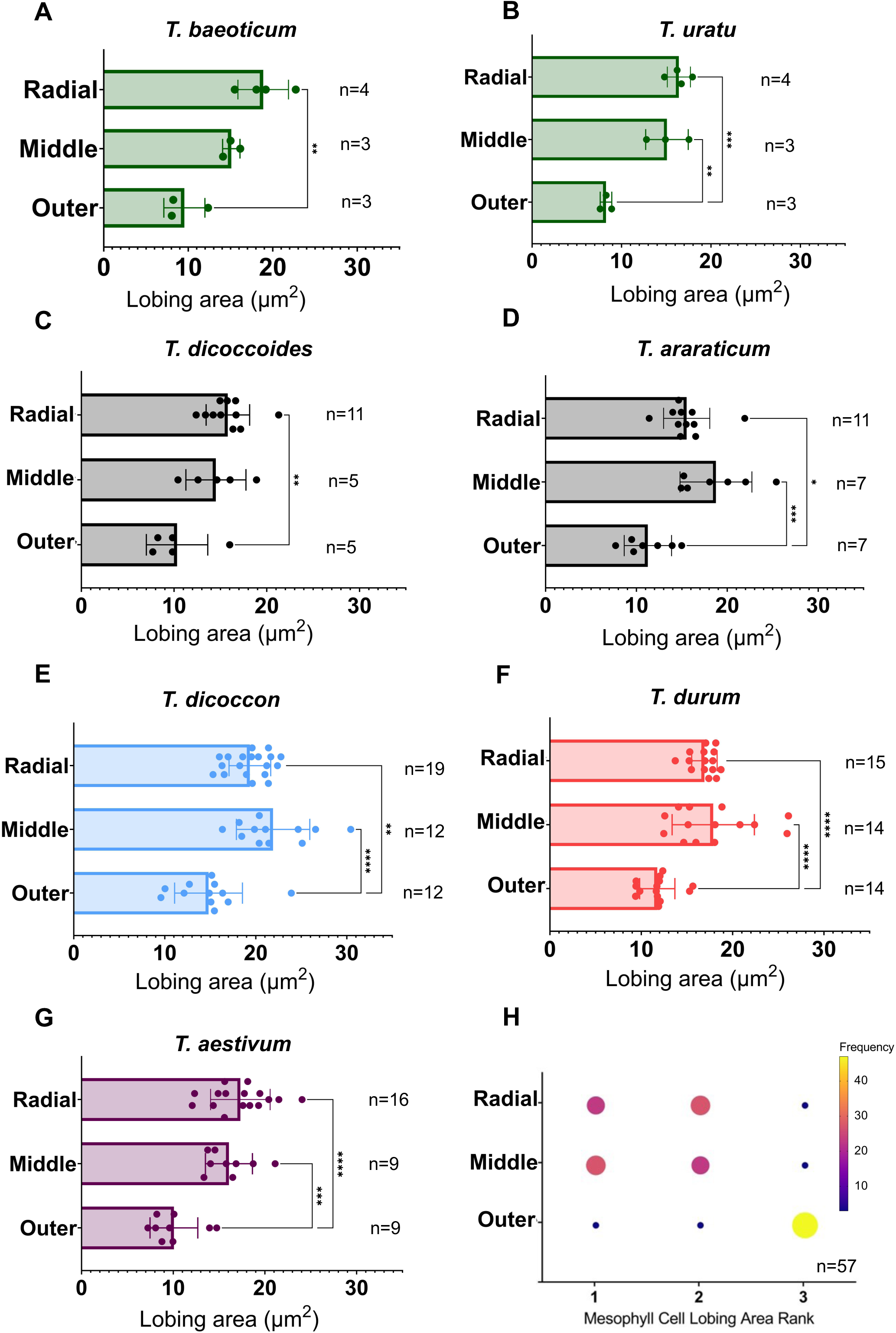
Analysis of longitudinal cell lobing area shows layers can be distinguished by shape in diploid, tetraploid and hexaploid Triticum lines. **A-B**) Longitudinal cell lobing area in designated mesophyll layers in diploid species, *T.baeoticum* and *T.uratu* (as indicated); **C-D**) wild tetraploid species *T. dicoccoides* and *T. araraticum*, as indicated. **E-F**) cultivated tetraploid species, *T. dicoccon* and *T. durum*, as indicated. **G**) hexaploid species, *T. aestivum*. **H**) Ranked data analysis of longitudinal mesophyll cell lobing area from all lines, separated by layer. Frequency of rank is indicated by colour and size of circle. Ranking is determined based on the frequency that cells in a particular layer rank 1st (most circular) – 3rd (least circular) in terms of cell circularity. Error bars represent standard deviation from the mean. ANOVA and post-hoc Tukey tests were carried out to test for significant interactions between cell layers within each species/subspecies: p= <0.05, p= <0.005, p= <0.0005, p= <0.0001. Statistical significance for ranked data in (H) was assessed using a Friedman test.

**Figure 5.**
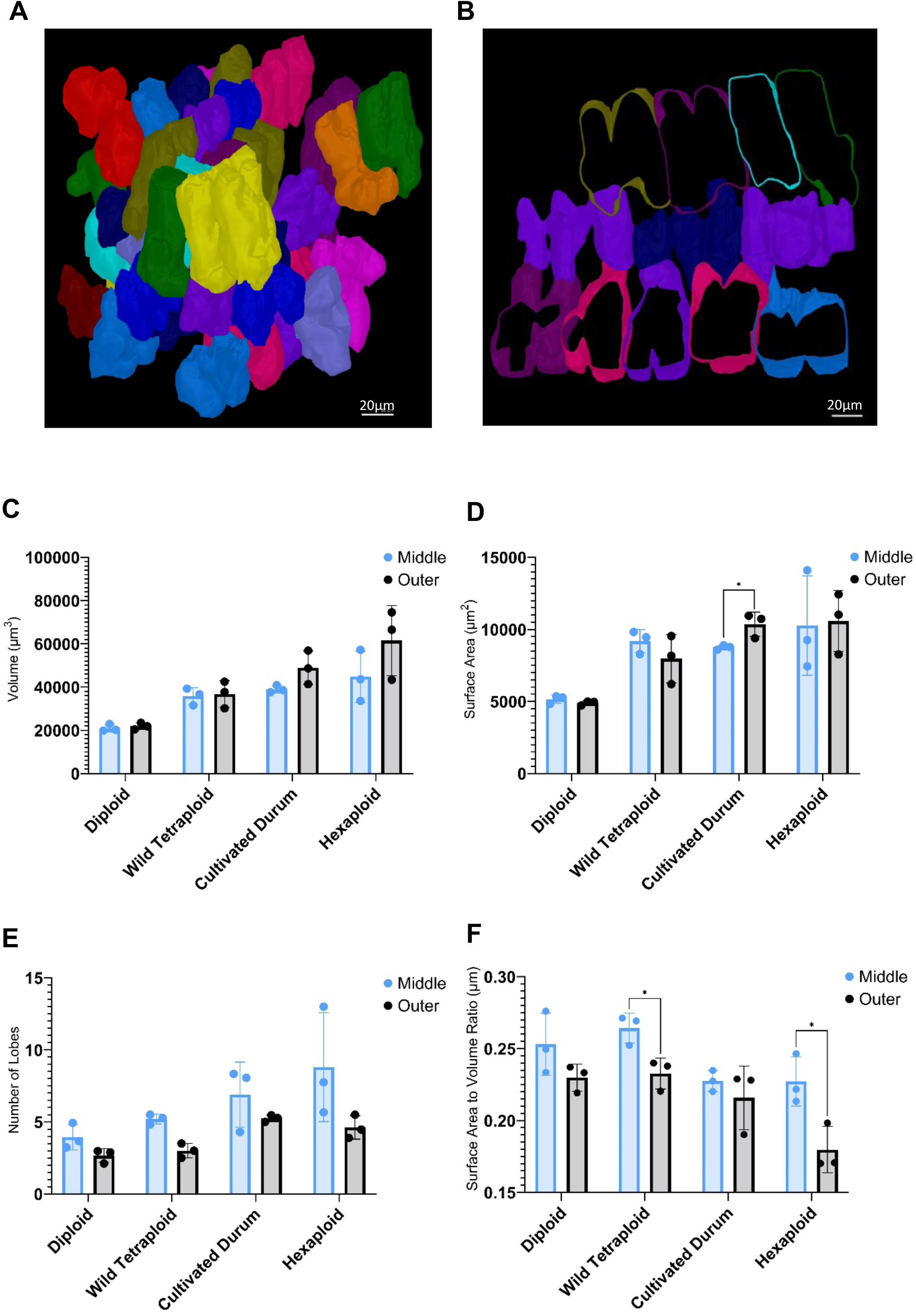
3D imaging and analysis of mesophyll cell volume, surface area and lobing in diploid, tetraploid and hexaploid Triticum reveals differences between layers. **A**) Portion of mesophyll of a wheat leaf (tetraploid T. *durum*, c.v. Voilur) after confocal imaging and segmentation to reveal mesophyll size and shape**. B**) Image showing clipped 3D cell objects generated from confocal stacks to reveal cell perimeter shape. From each 3D stack, ‘Outer’ and ‘Middle’ cells can be observed. **C**) Mesophyll cell volume in outer and middle mesophyll layers of diploid, tetraploid and hexaploid lines. **D**) Mesophyll cell surface area in outer and middle mesophyll layers of diploid, tetraploid and hexaploid lines. **E**) Number of lobes per cell in outer and middle mesophyll layers in diploid, tetraploid and hexaploid lines. **F**) Surface area to volume ratio of cells in outer and middle mesophyll layers in diploid, tetraploid and hexaploid lines. Error bars represent standard deviation from the mean. T-tests (A,B, D) and Mann-Whitney tests (C) were carried out to test for significant interactions between cell layers within each species/subspecies: p= <0.05, n=3.

With respect to cross-sectional cell area in transverse leaf sections, cells in the Middle and Radial layers had significantly smaller transverse mesophyll cell areas than those in Upper and Lower layers for all species/cultivars (**Figure 2**). The p values for the comparisons made (ANOVA followed by Tukey test) are indicated on the graphs, with differences between Middle/Radial vs Upper/Lower layers consistently being in a range <0.0001. Cell transverse areas of Middle and Radial layer mesophyll cells were indistinguishable from each other, and Upper and Lower layers could also not be distinguished from each other based on cell area, irrespective of ploidy level and cultivation status. There was an increase in mean mesophyll cell size with ploidy level (compare **Fig. 2A,B** (diploid) with **Fig. 2C-F** (tetraploid) and **Fig. 2G** (hexaploid), as has previously been reported (Wilson *et al,* 2021), with the patterns of cell size difference across the adaxial/abaxial axis superimposed on this shift in absolute cell size.

The patterning apparent in **Figure 2A-G** was strongly supported by a ranked data analysis (**Figure 2H**). In each leaf, the layers were assigned a rank of 1 to 4 based on the mean transverse mesophyll cell area. All plants from each ploidy level and cultivation status were grouped together and plotted using a heat map and circle size to represent the frequency of each layer having a specific rank. As demonstrated, the largest circles in the 1^st^ and 2^nd^ rank are always in the Upper and Lower mesophyll layers, with Middle and Radial cells the smallest (Friedman: p= <0.0001, Friedman statistic = 112.0). Statistically, the Upper and Lower layers both ranked lower than Radial and Middle cells at a 0.01% confidence interval (Dunn’s: p= <0.0001 for all interactions except Middle-Radial and Upper-Lower). The most common order from largest to smallest cells in the mesophyll across all ploidy levels was: Upper; Lower; Radial; Middle.

In contrast, a similar comparison of cell size according to the layers apparent in longitudinal leaf sections did not reveal major consistent differences in cell area (**Supplementary** Figure 1). The Outer and Middle layers could be distinguished in tetraploid *T. durum* and hexaploid *T. aestivum,* but this was not the case for other ploidy levels or cultivation status.

To investigate whether the differences in cross-sectional cell area in different layers revealed in **Figure 2** were reflected in variability of cell shape, we performed an analysis of cell circularity. These data showed that mesophyll layers differed significantly from each other in terms of cell circularity across all ploidy levels and cultivation status (**Figure 3**) (ANOVA p <0.05). The degree to which mesophyll cells within individual layers differed in circularity from other layers showed some variation depending on ploidy/cultivation status. For example, the diploid species were characterised by Middle/Radial cells being more circular than cells in the Upper/Lower layers (**Figure 3A,B**), largely mirroring the pattern observed in cell area (**Figure 2A,B**). In contrast, in wild tetraploid *T. araraticum,* cells in each layer could be distinguished from cells every other layer based on ci rcularity (**Figure 3D**), with Upper cells being the least circular, followed by Lower cells then Radial, and mesophyll cells in the Middle layer being the most circular.

From a combined ranked analysis (**Figure 3H**), the layer most frequently in the 1^st^ rank (most circular mesophyll cells) was the Middle layer, followed by Radial, Lower and Upper layers. This ranking was less well defined than for analysis based on transverse cell area (**Figure 2H**) but was still statistically significant (Friedman: p= <0.0001, Friedman statistic = 70.4), with the most consistent observation being that cells in the Upper layer were most commonly the least circular. The Upper-Middle, Upper-Radial and Upper-Lower rank differences were all statistically significant (Dunn’s: p= <0.0001; p= <0.0001; p= 0.0026 respectively). Cells in the Middle mesophyll layer were also consistently ranked higher (i.e. more circular) than both the Radial (Dunn’s: p= 0.0396) and Lower (Dunn’s: p= <0.0001), and cells in the Radial layer were indistinguishable from Lower cells in terms of transverse circularity.

Circularity gives a measure of the deviation of a shape from a perfect circle. Since it is apparent that the mesophyll cells tend to be lobed parallel to the leaf veins and there is substantial evidence that the change in surface area/volume is likely to influence parameters related to diffusion and, thus, mesophyll function, we also compared this parameter between cells in different layers. This was done using a “lobing area” function which measures the difference between the convex hull area of an object and the actual area that object (see Methods). Lobing in wheat mesophyll cells is most apparent in longitudinal sections of the leaf (as shown in **Figure 1**). Therefore, lobing area was calculated in layers defined in the longitudinal axis (Radial, Middle and Outer) (**Figure 4**). Differences in cell lobing area between layers were significant for all lines studied (*T. baeoticum* (ANOVA: p= 0.0048, f= 12.81), *T. uratu* (ANOVA: p= 0.0006, f= 25.22), *T. dicoccoides* (ANOVA: p= 0.0069, f= 6.646), *T. araraticum* (ANOVA: p= 0.0005, f= 10.94), *T. dicoccon* (ANOVA: p= <0.0001, f= 14.66), *T. durum* (ANOVA: p= <0.0001, f= 18.07) and *T. aestivum* (ANOVA: p= <0.0001, f= 18.42). Across all ploidy levels and cultivation statuses, Radial cells had significantly greater lobing area than Outer cells. Middle cells also had greater lobing area compared to cells in the Outer layers, with the exception of *T. baeoticum* (**Figure 4A**) and *T. dicoccoides* (**Figure 4D**), cells in Middle and Radial layers could not be distinguished from each other in terms of lobing area for any line.

When the data were combined and each layer ranked by lobing area, the observation of distinct layers was supported (Friedman: p= <0.0001, Friedman statistic: 54.94; **Figure 4H**). Cells in the Outer layers were ranked as consistently having the lowest lobing area compared to both Middle and Radial cells (Dunn’s: p= <0.0001 for both), with ranking of lobing area of the cells in the Middle and Radial layers not significantly different, i.e., they could not be distinguished from each other.

### Analysis of 3D mesophyll cell size and shape indicates differences across the adaxial/abaxial axis

Although measurements of cell size and shape in 2D sections of tissue are highly informative, cells are clearly 3D in nature. To obtain values of cell volume, surface area and shape, we performed a series of confocal imaging analyses, followed by cell segmentation in leaves from different species and cultivars of different ploidy and cultivation status. Z-stacks were acquired between the vasculature (i.e. omitting cells in the Radial layer), thus allowing for cells to be assigned to either Outer or Middle layers. Exemplar images are shown in **Figure 5A, B**, with quantitative data shown in **Figure 5C-F**. When cell volume was calculated, there was an increase in volume from diploid through to hexaploid, as expected (**Figure 5A**; ANOVA: p= 0.0009, f= 13.15). This increase in mesophyll cell volume with ploidy reflects an increase in cross-sectional area rather than cell length (**Figure 2**; **Supplementary** Figure 1). Comparison of Middle and Outer cells showed a trend for a relative increase in the volume of Outer mesophyll cells in cultivated tetraploid and hexaploid leaves, but this was not statistically significant (**Figure 5C**). In terms of mesophyll cell surface area, the only statistically significant difference was in *T. durum*, where the surface area of the Outer layer cells was greater than the Middle cells (t-test: p= 0.0318, t= 3.237, df= 4; **Figure 5D**).

With respect to lobe number, cells in the Middle layer had a higher number of lobes than the Outer cells at all ploidy levels and cultivation status (**Figure 5E**). When surface area/volume ratio was calculated (**Figure 5F**), mesophyll cells in the Middle layers had a greater surface are to volume ratio than Outer cells, which was statistically significant for both wild tetraploid (t-test: p= 0.022, t= 3.640, df= 4) and hexaploid lines (t-test: p= 0.025, t= 3.495, df= 4).

## DISCUSSION

We recently reported that rice leaves are characterised by a mesophyll in which there is a clear patterning of cell size and shape, with cells in the inner part of the leaf (not contacting the epidermis) characterised by being larger and more lobed than mesophyll cells adjoining the adaxial or abaxial epidermis of the leaf (Sloan *et al*., 2023). To investigate whether this patterning of the mesophyll was unique to rice or reflected a more general phenomenon in grass/cereal species, we performed a similar analysis on wheat leaves. Our data show that there is also a spatial pattern in mesophyll cell size and shape in wheat and that, moreover, this appears to have been conserved throughout selection/breeding of modern hexaploid bread wheat. The basal pattern is observed in extant diploid, tetraploid and hexaploid Triticum species, encompassing both wild and cultivated lines (**Figure 2, 3, 4**). The nature of this pattern is summarised schematically in **Figure 6**. In terms of size, the outer mesophyll cells are larger than the middle layer cells for all wheat species. This pattern of cell size (outer/middle) is maintained across species, despite the general increase in size of all cells with increase in ploidy (as has been previously reported, Wilson *et al*., 2021). There is also a pattern of lobing in which the middle layer cells are generally more lobed than the outer mesophyll cells across all species. The resultant adaxial/abaxial pattern leads to the mesophyll cells immediately sub-adjacent to the epidermis having an axiality which is generally perpendicular to the leaf surface, whereas the Middle layer cells are more highly lobed and have a long axis oriented generally parallel to the leaf surface.

**Figure 6.**
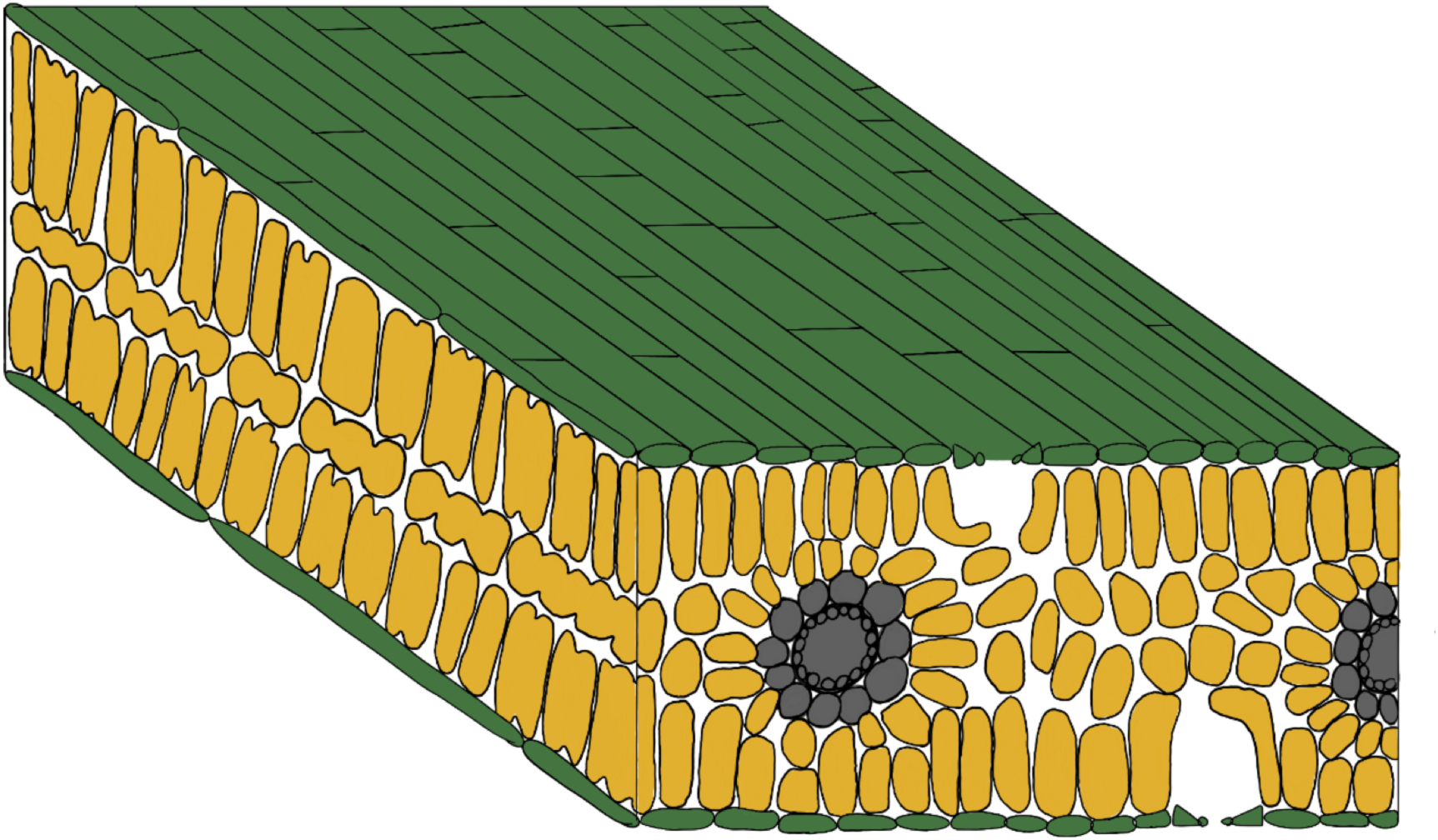
Summary schematic of typical triticale leaf structure. Mesophyll cells (orange) lie between upper and lower epidermis (green). The middle layer is distinguished by cells which are small but more lobed than upper and lower mesophyll cells adjoining the epidermis.

The functional significance of this spatial pattern of mesophyll cell size and shape in wheat awaits further elucidation. Eudicot leaves are characterised by a palisade layer in which the component cells have a cylindrical shape perpendicular to the leaf surface which plays an important role in initial light absorption (Parkhurst & Mott, 1990; Ho *et al*., 2016; Théroux-Rancourt & Gilbert, 2017; Borsuk & Brodersen, 2019). In particular, this mesophyll cell shape is thought to enable the focussing of light to inner cell layers in the eudicot leaf, helping to optimise light absorption and dispersal within the leaf (Vogelmann & Martin, 1993). Whether the outer mesophyll cells in wheat play a similar role awaits further investigation, but the challenge of maximising/optimising light distribution throughout the internal leaf structure is one that is likely to be common to all leaves. It has also been proposed that amphistomatous, isobilateral leaves – such as those showcased in the Triticeae – enable short diffusion pathways for CO_2_ by minimising the distance between stomata and the mesophyll (Drake *et al.,* 2019). The conservation of a high degree of lobing in the middle mesophyll cells might reflect a physical requirement linked to effective cell packing, as required for optimal CO_2_ flux. Alternatively, mesophyll cell lobing has long been linked to providing a relatively large surface area for gas exchange in addition to increasing the chloroplast area exposed to intercellular airspaces (Sage & Sage, 2009), and is therefore thought to be an important parameter in gas diffusion underpinning CO_2_ absorption for photosynthesis (Terashima *et al.,* 2011; Ren *et al.,* 2019). Thus, it is possible that the relatively high degree of lobing in the middle mesophyll cells of wheat enables more effective gas exchange in these cells.

Although the patterns described above were observed in all wheat species, there were differences between the species investigated, most notably between the cultivated and non-cultivated lines. For example, when cell volume was measured, there was a trend for increased size of the Outer-layer mesophyll cells relative to the Middle-layer cells in the cultivated Triticum varieties compared to non-cultivated, i.e., mesophyll cell volume was more uniform in the non-cultivated species. This is obviously based on the observation of a limited set of species, but does raise the question of whether during breeding there has been an unintentional selection for a leaf architecture of larger outer mesophyll cells compared with the Middle-layer. If this is the case, this raises the question of whether this has had a functional consequence for the physiological performance of leaves in cultivated wheat.

Our finding of a conserved pattern of mesophyll cell size and shape in wheat adds to our report of a similar (yet distinct) pattern in rice (Sloan *et al.,* 2023). In particular, in wheat the middle layer cells are relatively small, whereas in rice the middle layer cells are relatively large. The functional significance of this difference is unclear. Overall, our data support the hypothesis that there is an ancient conserved spatial pattern of cells in the grass mesophyll, and also show that shifts in the balance of relative size/shape of cells in different layers can occur during breeding/selection. Our results provide a platform for future studies to investigate the functional significance of grass mesophyll patterning at cellular resolution.

## Acknowledgements

E.W. was supported by a BBSRC-White Rose DTP (BB/M011151) CASE studentship with RAGT Seeds Ltd (R.S.) to A.J.F. and C.U.. M.J.W. was supported by BBSRC grants ‘‘Shape Shifting Stomata: The Role of Geometry in Plant Cell Function’’ (BB/T005041) and “A New Model of Stomatal Function” (BB/Y001257/1) to A.J.F.. Imaging was performed at the Wolfson Light Microscopy Facility at the University of Sheffield

## Contributions

Conceptualisation: A.J.F, C.U.; investigation: E.W., M.J.W..; writing – original draft, E.W., A.J.F..; writing – review & editing: all authors; supervision: A.J.F., M.J.W., R.S., C.U.; funding acquisition: A.J.F., C.U., R.S.

## SUPPLEMENTARY INFORMATION

**Supplementary Fig. 1.**
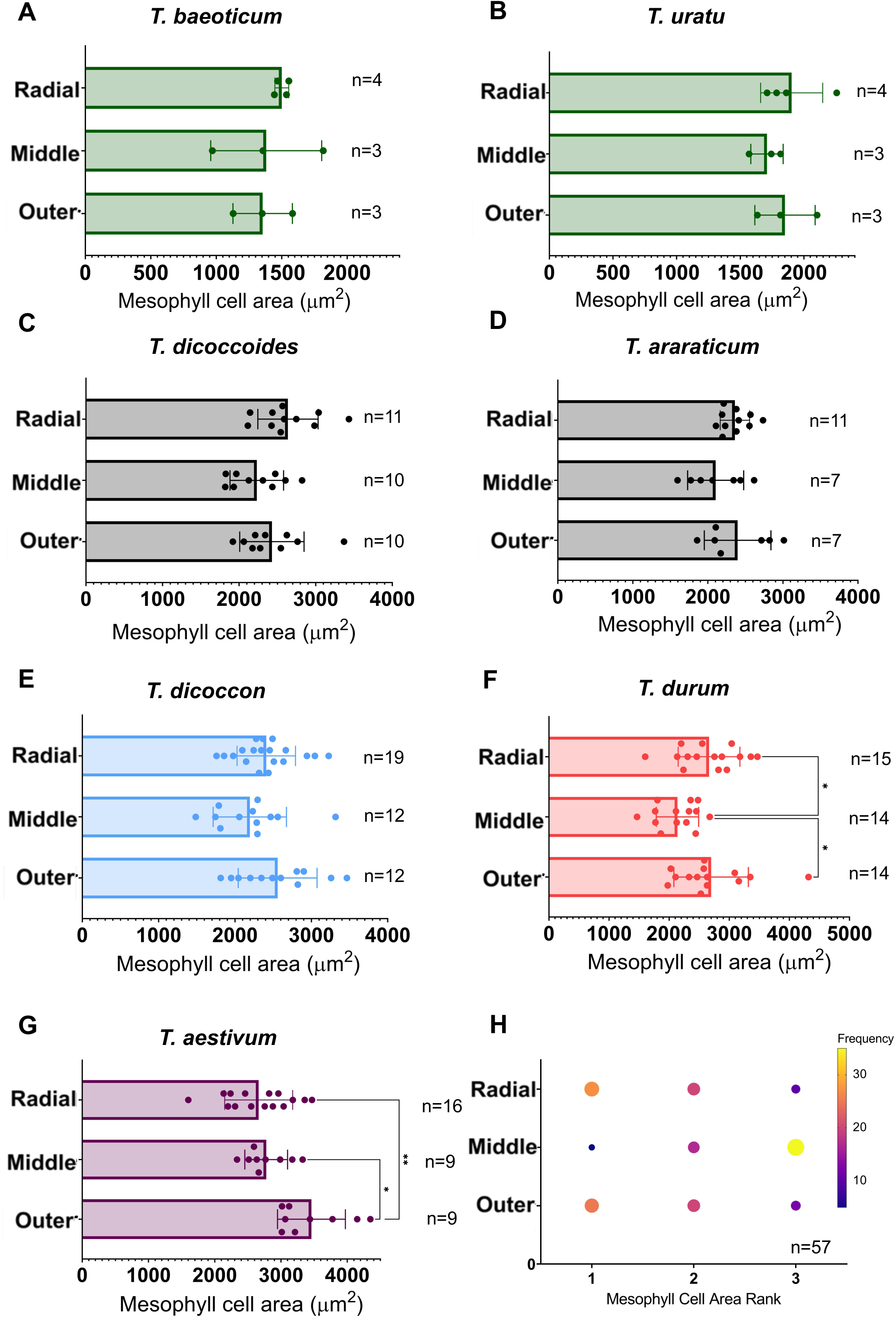
Analysis of longitudinal mesophyll cell area shows layers cannot be consistently distinguished by size in diploid, tetraploid and hexaploid Triticum lines. **A-B**) Longitudinal area of cells in designated mesophyll layers of diploid species, *T.baeoticum* and *T.uratu* (as indicated); **C-D**) tetraploid species, *T. dicoccoides* and *T. araraticum*, as indicated; **E-F**) cultivated tetraploid species, *T. dicoccon* and *T. durum*, as indicated; **G**) hexaploid species, *T. aestivum*; **H**) Ranked data of longitudinal mesophyll cell area from all lines, separated by layer. Frequency of rank is indicated by colour and size of circle. Ranking is determined based on the frequency that cells in a particular layer rank 1st (largest) –4th (smallest) in terms of cell area. In A-G, error bars represent standard deviation from the mean. ANOVA and post-hoc Tukey tests were carried out to test for significant interactions between cell layers within each species/subspecies: p= <0.005, p= <0.0005, p= <0.0001. Statistical significance for ranked data in (H) was assessed using a Friedman test.

## Notes

### Competing Interest Statement

The authors have declared no competing interest.

